# Quantifying the turnover of transcriptional subclasses of HIV-1-infected cells

**DOI:** 10.1101/000778

**Authors:** Christian L. Althaus, Beda Joos, Alan S. Perelson, Huldrych F. Günthard

**Affiliations:** Institute of Social and Preventive Medicine (ISPM), University of Bern, 3012 Bern, Switzerland; Division of Infectious Diseases and Hospital Epidemiology, University Hospital Zürich, University of Zürich, Zürich, Switzerland; Theoretical Division, Los Alamos National Laboratory, Los Alamos, New Mexico 87545, United States of America

## Abstract

HIV-1-infected cells in peripheral blood can be grouped into different transcriptional subclasses. Quantifying the turnover of these cellular subclasses can provide important insights into the viral life cycle and the generation and maintenance of latently infected cells. We used previously published data from five patients chronically infected with HIV-1 that initiated combination antiretroviral therapy (cART). Patient-matched PCR for unspliced and multiply spliced viral RNAs combined with limiting dilution analysis provided measurements of transcriptional profiles at the single cell level. Furthermore, measurement of intracellular transcripts and extracellular virion-enclosed HIV-1 RNA allowed us to distinguish productive from non-productive cells. We developed a mathematical model describing the dynamics of plasma virus and the transcriptional subclasses of HIV-1-infected cells. Fitting the model to the data allowed us to better understand the phenotype of different transcriptional subclasses and their contribution to the overall turnover of HIV-1 before and during cART. The average number of virus-producing cells in peripheral blood is small during chronic infection (25.7 cells ml^−1^). We find that 14.0%, 0.3% and 21.2% of infected cells become defectively, latently and persistently infected cells, respectively. Assuming that the infection is homogenous throughout the body, we estimate an average *in vivo* viral burst size of 2.1 × 10^4^ virions per cell. Our study provides novel quantitative insights into the turnover and development of different subclasses of HIV-1-infected cells. The model predicts that the pool of latently infected cells becomes rapidly established during the first months of acute infection and continues to increase slowly during the first years of chronic infection. Having a detailed understanding of this process will be useful for the evaluation of viral eradication strategies that aim to deplete the latent reservoir of HIV-1.

**Author Summary:** Gaining a quantitative understanding of the development and turnover of different HIV-1-infected subpopulations of cells is crucial to improve the outcome of patients on combination antiretroviral therapy (cART). The population of latently infected cells is of particular interest as they represent the major barrier to a cure of HIV-1 infection. We developed a mathematical model that describes the dynamics of different transcriptionally active subclasses of HIV-1-infected cells and the viral load in peripheral blood. The model was fitted to previously published data from five chronically HIV-1-infected patients starting cART. This allowed us to estimate critical parameters of the within-host dynamics of HIV-1, such as the the number of virions produced by a single infected cell. The model further allowed investigation of HIV-1 dynamics during the acute phase. Computer simulations predict that latently infected cells become rapidly established during the first months of acute infection and continue to increase slowly during the first years of chronic infection. This illustrates the opportunity for strategies that aim to eradicate the virus during early cART as the pool of HIV-1 infected cells is substantially smaller during acute infection than during chronic infection.

## Introduction

High levels of cell-associated HIV-1 RNA can be observed in peripheral blood of patients with undetectable plasma viremia during combination antiretroviral therapy (cART) [1–4]. The various HIV-1 RNA and DNA species that are present during the viral life cycle can serve as biomarkers for basal transcription in viral reservoirs with different properties [5, 6]. Gaining a quantitative understanding of the development and turnover of HIV-1-infected subpopulations and viral latency is of particular interest in light of recent efforts in viral eradication strategies [7–10].

Highly sensitive assays for HIV-1 plasma RNA in patients on cART usually provide bulk measurements of viral activity and cannot distinguish between different infected subpopulations [11]. In contrast, the study by Fischer et al. [12] combined highly sensitive PCR assays for unspliced (UsRNA) and multiply spliced (MsRNA-tatrev and MsRNA-nef) HIV-1 RNA species with limiting dilution endpoint analysis of peripheral blood mononuclear cells (PBMCs). In addition to intracellular RNA transcripts, extracellular virion-enclosed HIV-1 RNA that provides a marker for cells releasing virus particles was also measured. The study identified four distinct viral transcriptional classes: two overlapping cell classes of high viral transcriptional activity, representative of a virus producing phenotype; and two cell classes that express HIV-1 RNA at low and intermediate levels that match definitions of viral latency [12, 13].

Analyzing the decay kinetics of plasma viral load in HIV-1-infected patients on cART using mathematical models has resulted in a detailed understanding of viral replication dynamics *in vivo* [14–16]. The plasma viral load typically exhibits three exponential phases during the first year after start of cART (Figure 1). Due to the rapid turnover of free virus in blood [17], the viral decay phases are thought to reflect the contribution of different HIV-1-infected cell populations on viral production. The first phase with a half-life of 1 to 2 days is attributed to the loss of activated, virus-producing cells [18, 19]. The second phase exhibits a half-life of 1 to 4 weeks and is considered to reflect the loss of so-called persistently infected cells such as resting CD4^+^ T cells or macrophages [20]. Finally, the third phase decay has a long half-live of 39 weeks suggesting that latently infected cells are a primary candidate for this cellular compartment [21, 22], although slow release of virus from the follicular dendritic cell network is another possibility [23]. Other mathematical models have been developed that stratify the infected cells into additional subpopulations such as non-productively infected cells during the intracellular eclipse phase [24] and defectively infected cells [25]. Nevertheless, most studies to date are focused on the analysis of viral load and only indirectly allow inferring the kinetics of cellular subpopulations. Few studies have attempted to characterize the concentration of virus and several infected subpopulations based on data simultaneously [25]. Fitting mathematical models to multiple quantities of viral replication would result in refined parameter estimates for describing the generation and maintenance of latently infected cells.

**Figure 1.**
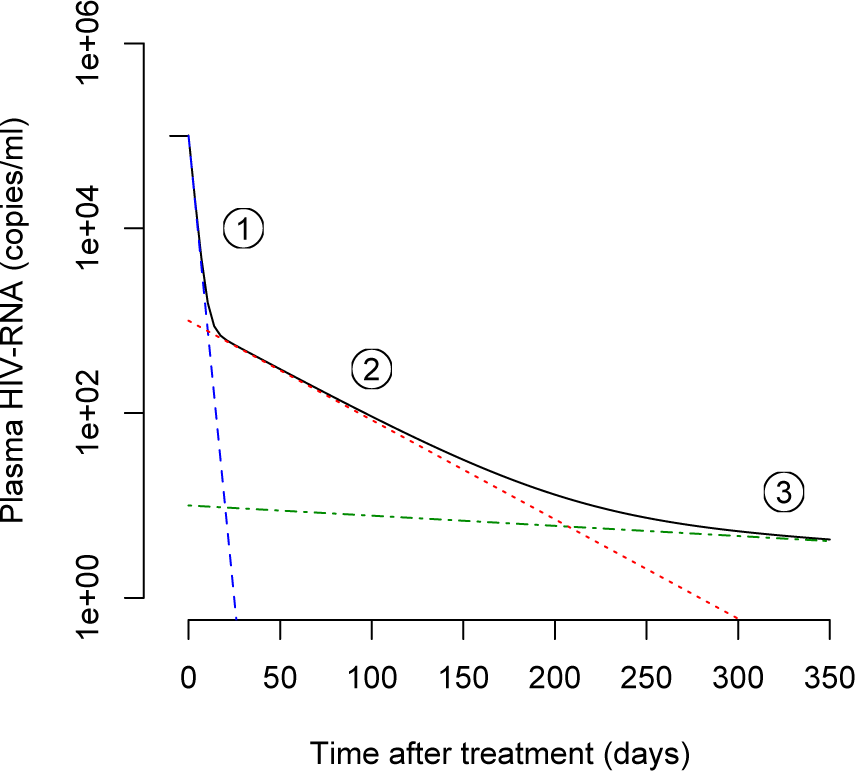
Three-phase decay of HIV-1 after start of cART. The black line shows a typical decay profile of plasma viral load during the first year of cART. Typical half-lives of the first (blue dashed line), second (red dotted line) and third phase (green dash-dotted line) are 1.5 days, 4 weeks and 39 weeks, respectively [21].

In this study, we developed a mathematical model that describes the dynamics of different transcriptionally active subclasses of HIV-1-infected cells and the viral load in peripheral blood. The model was fitted to previously published data from five chronically HIV-1-infected patients starting cART [12]. This allowed us to estimate critical parameters of the within-host dynamics of HIV-1 and the turnover of various subpopulations of infected cells. Finally, we simulated the development of the latently infected cell pool during acute infection, providing useful information for viral eradication strategies.

## Results

We first devised a detailed model of the within-host dynamics of HIV-1 that is based on the observations of different subclasses of HIV-1-infected cells in the study by Fischer et al. [12]. The five subclasses are HIV-1 DNA^+^, low, medium and high HIV-1 RNA expressing and cells that have virion-enclosed HIV-1 RNA associated with them (also see *Methods*). These subclasses show distinct decay dynamics in patients on cART (Figure 2). The slow decay of the subclass of PBMCs that contains proviral DNA (*DNA*^+^) indicates that this cell population primarily contributes to the third phase decay and likely consists of defectively or latently infected cells to a large extent. The subclasses of PBMCs that contain viral RNA all show a rapid drop early after start of cART. We hypothesize that this is due to the loss of different subpopulations of actively infected cells that rapidly move through the intracellular eclipse phase before they start to produce virus particles and die. After the early drop, the subclass of cells exhibiting UsRNA only (*Low)* decays slowly and most likely consists mainly of latently infected cells with low basal transcription of HIV-1. The cells with medium transcriptional activity (*Mid)* appear to contribute to the second and the third phase viral decay, which is characteristic of persistently and latently infected cells. The early drop in PBMCs with a higher transcriptional activity (*High*), which is more pronounced compared to cells with a low and medium transcriptional activity, that is followed by a slower loss of cells is reminiscent of activated, virus-producing and persistently infected cells. Finally, the PBMCs that have extracellular virion-enclosed HIV-1 RNA associated with them (*Extra*) show a very rapid loss before reaching the limit of detection. This is expected as they should represent the short-lived population of virus-producing cells [4] that contribute to the first phase of viral decay.

**Figure 2.**
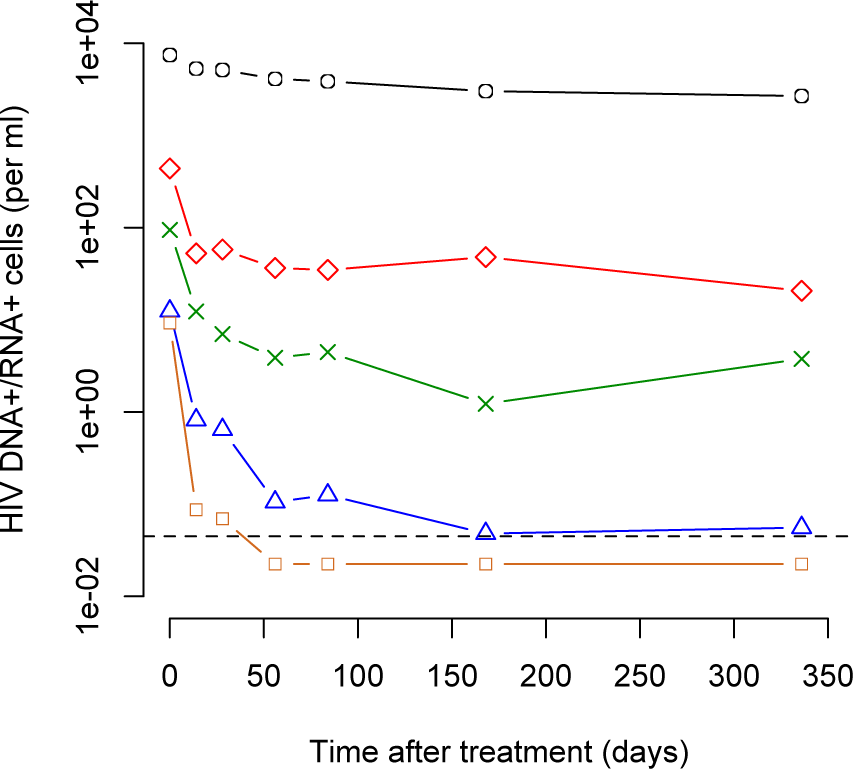
Decay kinetics of subclasses of HIV-1-infected cells during cART. The five subclasses of cells are: DNA^+^ (black circles), Low (red diamonds), Mid (green crosses), High (blue triangles) and Extra (chocolate squares). Symbols represent geometric means of the five patients from the study by Fischer et al. [12]. The dashed line represents the limit of detection that was set at 50% of the lowest measured cell count. Measurements below this threshold were assumed to be at 50% of the detection limit to include them in the mean.

The different subclasses of HIV-1-infected cells clearly overlap and are representative of heterogeneous cell populations. Furthermore, the life cycle of HIV-1 from infection of a cell to the release of virus particles can be divided into cell populations with different transcriptional activity [26]. We took both of these important characteristics into account in our model that consists of 12 subpopulations of cells that can be stratified according to their HIV-1 DNA and RNA content (Figure 3 and *Methods*). In this model, we defined persistently infected cells (*M*_1_ and *M*_2_) as long-lived cells that can produce viral particles. Latently infected cells (*L*_1_ and *L*_2_) were assumed to transcribe HIV-1 RNA at low or intermediate levels [12, 13]. Infected cells that are HIV-1 DNA positive, but HIV-1 RNA negative, were assumed to remain transcriptionally silent during the observation period and considered as defectively infected cells (*D*).

**Figure 3.**
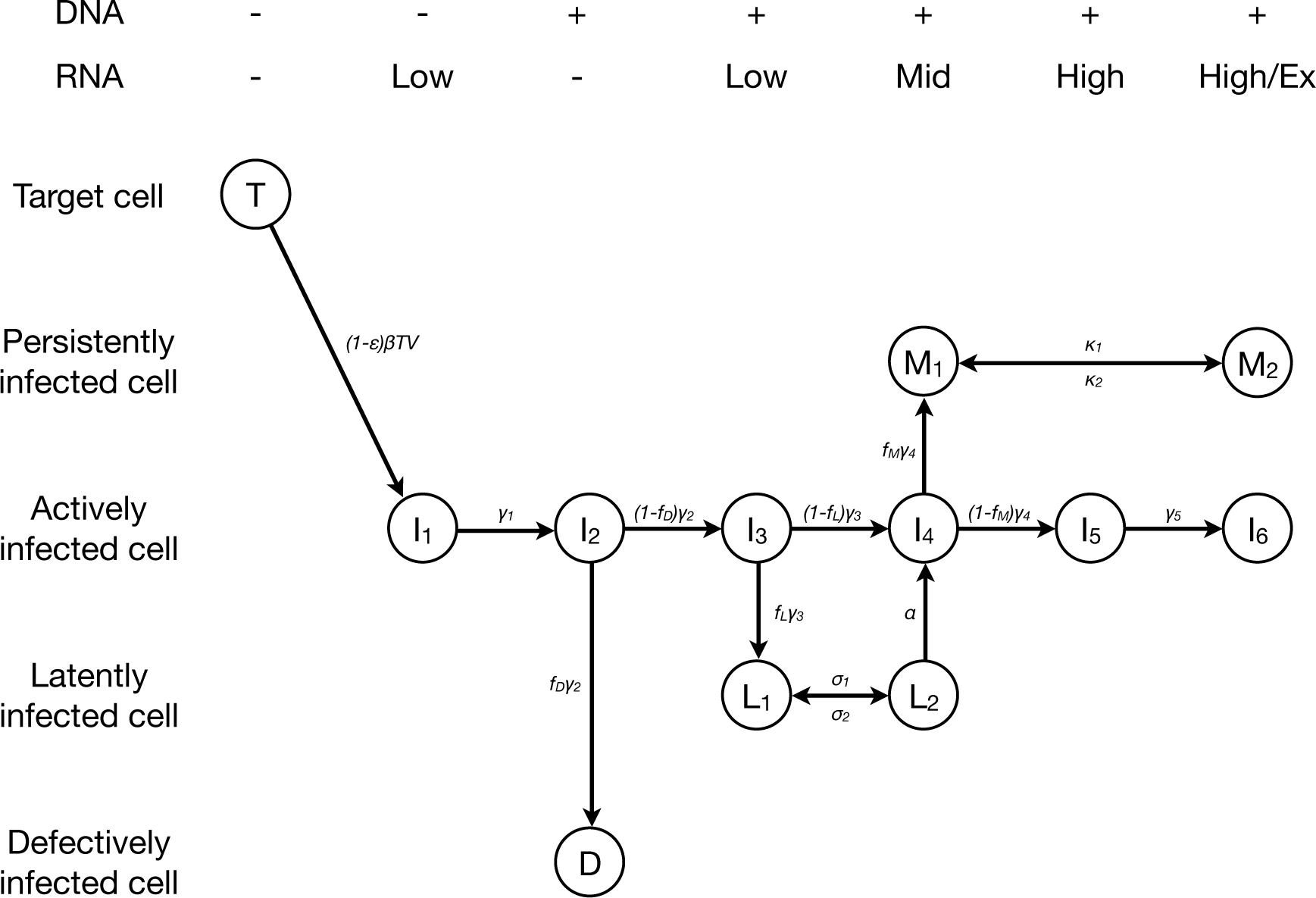
Model of HIV-1 dynamics. Actively infected cells move through an intracellular eclipse phase (*I*_1_ to *I*_5_) before they start to produce virus particles (*I*_6_). Some of the cells during the intracellular eclipse phase become either defectively infected (*D*), latently infected (*L*_1_) or persistently infected (*M*_1_). Both latently (*L*_1_ and *L*_2_) and persistently (*M*_1_ and *M*_2_) can move between two transcriptional states. Persistently infected cells that are in a high transcriptional state (*M*_2_) also contribute to virus production. The different subpopulations of infected cells can be stratified according to their HIV-1 DNA and RNA content (shown on top).

Fitting the mathematical model to the data from five HIV-1-infected patients resulted in a good description of the viral and cellular decay kinetics during cART (Figure 4 and *Text S1*). The model clearly describes more pronounced decay dynamics in infected cells with increasing transcriptional activity. Table 1 gives a summary of the best fit estimates of the model parameters that describe the virus dynamics. We found that 1.1% (0.2%–7.0%) of all CD4^+^ T cells can be target cells for infection with HIV-1. This is somewhat lower than the 6.5% of CD4^+^ Ki-67^+^ T cells in HIV-1-infected individuals that have been measured previously [27]. We also obtained estimates for the average lifespans of target cells (61 days, range: 11–528 days) and latently infected cells (33 years, range: 168 days–505 years). While others have estimated the average half-life of latently infected cells to be 6.3 months [28] and 44 months [29], our estimates are less precise due to the much shorter follow-up period after start of cART. However, the estimated activation rate of latently infected cells (2.7 *×* 10^−3^ d^−1^, range: 5.3 *×* 10^−5^ − 1.5 *×* 10^−1^ d^−1^) that also influences the slope of the third phase decay in plasma HIV-1 RNA is consistent with previous findings [30].

**Figure 4.**
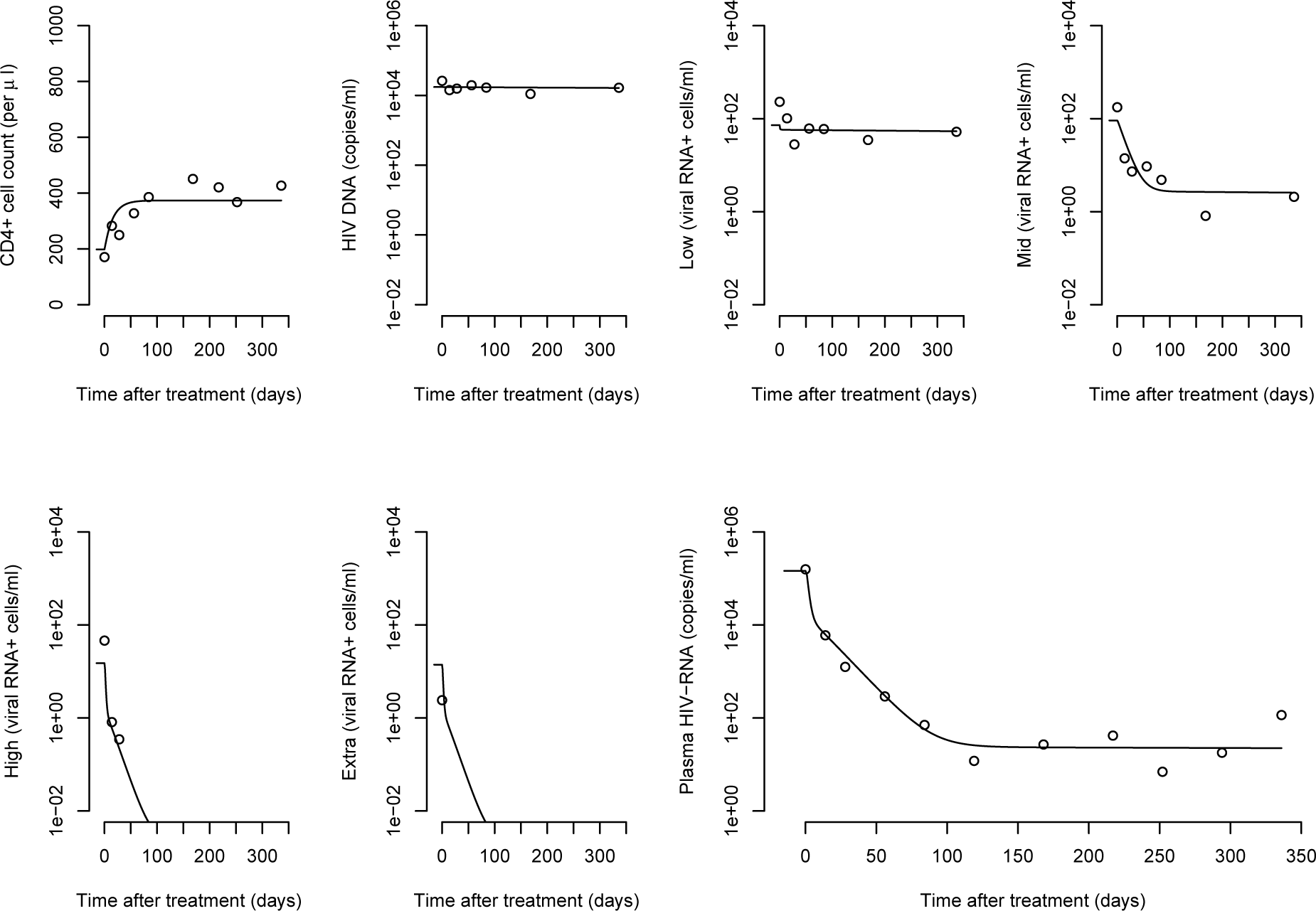
HIV-1 dynamics during cART. Circles denote measured data of patient 112 and lines represent the best fit of the default model. Model fits to data of the four other patients are given in *Text S1*.

**Table 1.**
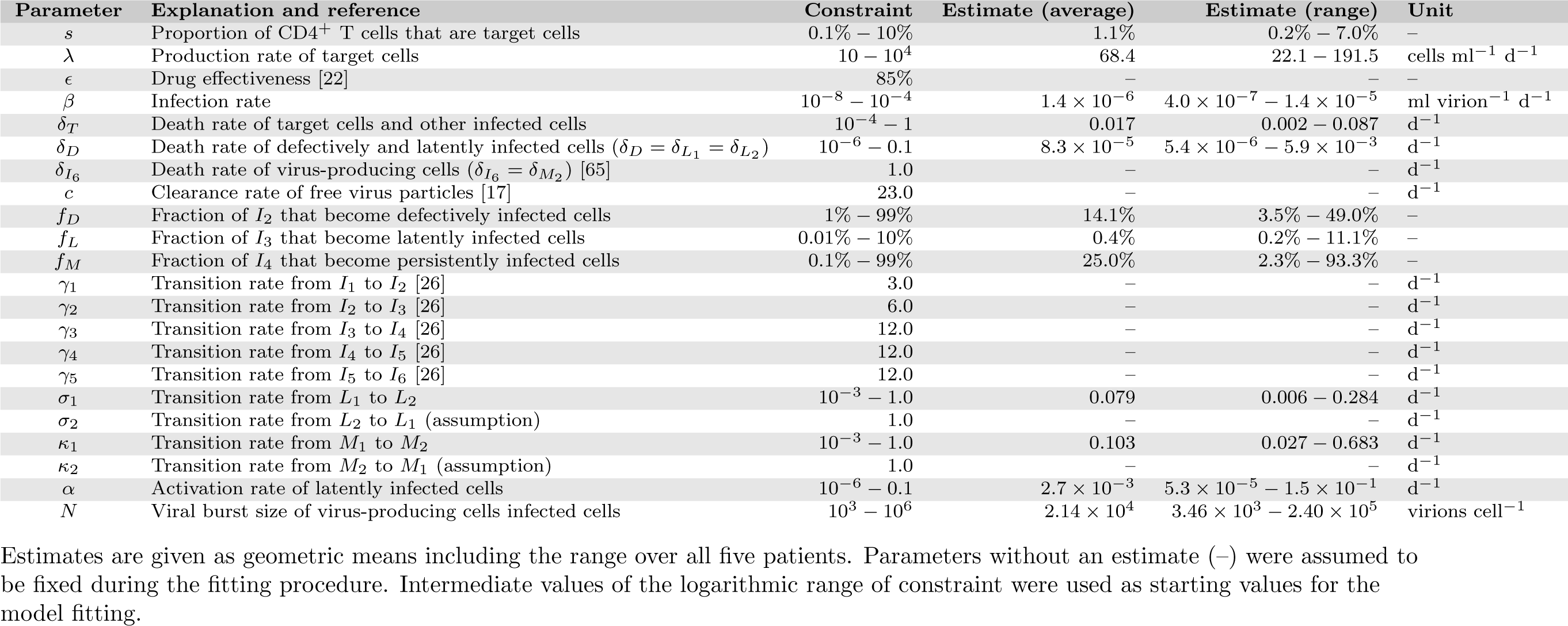
Estimated parameters of HIV-1 dynamics.

| Parameter | Explanation and reference | Constraint | Estimate (average) | Estimate (range) | Unit |
| --- | --- | --- | --- | --- | --- |
| $s$ | Proportion of CD4<sup>+</sup> T cells that are target cells | 0.1% – 10% | 1.1% | 0.2% – 7.0% | – |
| $\lambda$ | Production rate of target cells | $10 - 10^4$ | 68.4 | 22.1 – 191.5 | cells ml<sup>-1</sup> d<sup>-1</sup> |
| $\epsilon$ | Drug effectiveness [22] | 85% | – | – | – |
| $\beta$ | Infection rate | $10^{-8} - 10^{-4}$ | $1.4 \times 10^{-6}$ | $4.0 \times 10^{-7} - 1.4 \times 10^{-5}$ | ml virion<sup>-1</sup> d<sup>-1</sup> |
| $\delta_T$ | Death rate of target cells and other infected cells | $10^{-4} - 1$ | 0.017 | 0.002 – 0.087 | d<sup>-1</sup> |
| $\delta_D$ | Death rate of defectively and latently infected cells ($\delta_D = \delta_{L_1} = \delta_{L_2}$) | $10^{-6} - 0.1$ | $8.3 \times 10^{-5}$ | $5.4 \times 10^{-6} - 5.9 \times 10^{-3}$ | d<sup>-1</sup> |
| $\delta_{I_6}$ | Death rate of virus-producing cells ($\delta_{I_6} = \delta_{M_2}$) [65] | 1.0 | – | – | d<sup>-1</sup> |
| $c$ | Clearance rate of free virus particles [17] | 23.0 | – | – | d<sup>-1</sup> |
| $f_D$ | Fraction of $I_2$ that become defectively infected cells | 1% – 99% | 14.1% | 3.5% – 49.0% | – |
| $f_L$ | Fraction of $I_3$ that become latently infected cells | 0.01% – 10% | 0.4% | 0.2% – 11.1% | – |
| $f_M$ | Fraction of $I_4$ that become persistently infected cells | 0.1% – 99% | 25.0% | 2.3% – 93.3% | – |
| $\gamma_1$ | Transition rate from $I_1$ to $I_2$ [26] | 3.0 | – | – | d<sup>-1</sup> |
| $\gamma_2$ | Transition rate from $I_2$ to $I_3$ [26] | 6.0 | – | – | d<sup>-1</sup> |
| $\gamma_3$ | Transition rate from $I_3$ to $I_4$ [26] | 12.0 | – | – | d<sup>-1</sup> |
| $\gamma_4$ | Transition rate from $I_4$ to $I_5$ [26] | 12.0 | – | – | d<sup>-1</sup> |
| $\gamma_5$ | Transition rate from $I_5$ to $I_6$ [26] | 12.0 | – | – | d<sup>-1</sup> |
| $\sigma_1$ | Transition rate from $L_1$ to $L_2$ | $10^{-3} - 1.0$ | 0.079 | 0.006 – 0.284 | d<sup>-1</sup> |
| $\sigma_2$ | Transition rate from $L_2$ to $L_1$ (assumption) | 1.0 | – | – | d<sup>-1</sup> |
| $\kappa_1$ | Transition rate from $M_1$ to $M_2$ | $10^{-3} - 1.0$ | 0.103 | 0.027 – 0.683 | d<sup>-1</sup> |
| $\kappa_2$ | Transition rate from $M_2$ to $M_1$ (assumption) | 1.0 | – | – | d<sup>-1</sup> |
| $\alpha$ | Activation rate of latently infected cells | $10^{-6} - 0.1$ | $2.7 \times 10^{-3}$ | $5.3 \times 10^{-5} - 1.5 \times 10^{-1}$ | d<sup>-1</sup> |
| $N$ | Viral burst size of virus-producing cells infected cells | $10^3 - 10^6$ | $2.14 \times 10^4$ | $3.46 \times 10^3 - 2.40 \times 10^5$ | virions cell<sup>-1</sup> |

Estimates are given as geometric means including the range over all five patients. Parameters without an estimate (–) were assumed to be fixed during the fitting procedure. Intermediate values of the logarithmic range of constraint were used as starting values for the model fitting.

The parameters *f_D_*, *f_L_* and *f_M_* denote the fractions of cells that end up in a particular subpopulation in a sequential process during the intracellular eclipse phase. From this, we can calculate the average proportion of newly infected cells that become a certain cell type (*Text S1)*. In contrast to another study [25], we find that only 63.4% of infected cells become activated, virus-producing cells (*I*_6_). A substantial fraction of infected target cells results in defectively (14.0%) and persistently infected cells (21.2%). The proportion of infected cells that become latently infected or die before ending up in one of the subpopulations is small (0.3% and 1.1%, respectively). Note that after activation, latently infected cells can then either become persistently infected or activated, virus-producing cells by moving through cell class *I*_4_. Transcriptional bursts that increase the level of viral RNA transcription occur on average every 12.7 days (1*/σ*_2_, range: 3.5–165.2 days) and 9.7 days (1*/κ*_2_, range: 1.5–37.0 days) in latently and persistently infected cells, assuming that bursts last for one day on average (*σ*_1_ = *κ*_1_ = 1 d^−1^). The total number of virus particles produced by a cell during its lifetime, the viral burst size, was estimated at 21’387 virions per cell (range: 3459–239’770 virions per cell). Note that we assumed that persistently infected cells in an elevated transcriptional state (*M*_2_) produce viral particles at the same rate as activated, virus-producing cells. However, the duration of virus release is shorter in persistently infected cells as they can revert to a lower transcriptional state (*M*_1_). The high viral burst size suggests that the numbers of virus-producing cells in peripheral blood must be small and we indeed found an average of only 25.7 cells ml^−1^ (range: 7.8–143.1 cells ml^−1^) in the model during the chronic phase of infection.

The parameters were estimated by fitting the virus dynamics model to data of patients chronically infected with HIV-1. Based on those estimates, the model also allows investigation of virus dynamics during the acute phase. We used the average of the estimated parameters to simulate early infection with HIV-1 from a small viral inoculum in a hypothetical patient. We set *V* (0) = 1 copy per ml and assumed that the target cells are at steady-state (*T* (0) = *λ/δ_T_*). The rapid rise of plasma HIV-1 RNA during the first weeks of infection is followed by the chronic phase at which the virus concentration reaches its set-point level (Figure 5). The total pool of latently infected cells (*L*_1_ + *L*_2_) show somewhat different dynamics during acute HIV-1 infection. A very rapid expansion of latent cells during the viral growth phase is followed by a slower increase into the chronic phase of infection. From the time of peak viremia (22 days) to the chronic phase (1000 days), the latently infected cell pool expands 14.3-fold from 9.8 to 140.4 cells per ml. The expansion of the total number of HIV-1 DNA positive cells from the acute (1813 cells per ml) to the chronic phase (7608 cells per ml) is smaller (4.2-fold). This is consistent with the 3.8-fold difference in the number of HIV-1 DNA copies that were measured in patients that initiated cART during the acute and chronic phase from another study [31, and see *Text S1*]. The time after infection at which latently infected and HIV-1 DNA positive cells reach 50% of their chronic level is 441 and 451 days, respectively. Altogether, this illustrates the opportunity for eradication strategies during early cART interventions as the pool of HIV-1 infected cells is substantially smaller during acute infection than during chronic infection.

**Figure 5.**
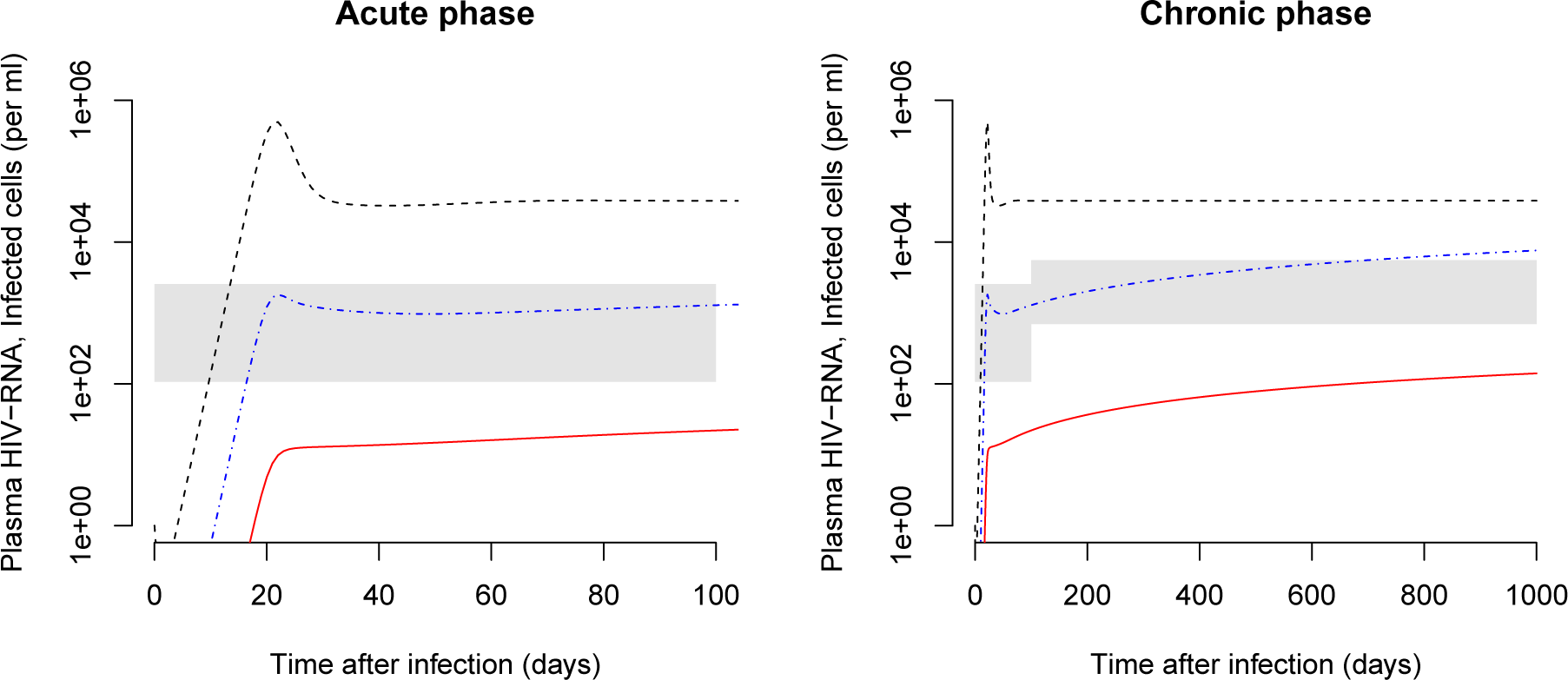
Development of the latently infected cell pool during acute and chronic of HIV-1 infection. The pool of latently infected cells (*L*_1_ + *L*_2_) is shown as a red line, the number of HIV-1 DNA positive cells as a blue dash-dotted line and the plasma HIV-1 RNA as a black dashed line. The average parameter estimates from Table 1 were used for the model simulation. The gray areas represent two standard deviations around the mean of the number of HIV-1 DNA positive cells in patients that initiated cART during acute and chronic infection [31] (for details, see *Text S1)*. The chronic phase graph depicts the same simulation as the acute phase graph but taken out to a longer time (1000 days).

## Discussion

We present the first mathematical model of virus dynamics that groups the different subpopulations of HIV-1-infected cells according to their transcriptional profile. The model assumes a heterogeneous population of latently and persistently infected cells having occasional transcriptional bursts to increase their level of RNA transcription which is consistent with experimental data from Fischer et al. [12]. Fitting this model to the unique data of virus transcription levels at the single cell level resulted in new estimates of the HIV-1 dynamics *in vivo*. We found that a large fraction of infected cells become either defectively or persistently infected cells. Furthermore, we found that the viral burst size can be high, between 3.5 *×* 10^3^ and 2.4 *×* 10^5^ viral particles per virus-producing cell. Lastly, we simulated the acute phase of HIV-1 infection in a typical patient. This illustrated that the latently infected cell pool becomes rapidly established during the first months of acute infection and shows a slow increase during the first years of chronic infection.

Our study is unique in that we fit a mathematical model of HIV-1 within a host to data of the virus dynamics at the single cell level. This is a substantial step beyond modeling studies that considered free virus in plasma, CD4^+^ T cells and bulk measurements of viral activity only. The new quantitative insights into the replication dynamics of HIV-1 *in vivo* that this study provides will be useful for an improved understanding of HIV and the effects of novel treatment strategies.

The measurements of HIV-1-infected cells and the virus concentration were performed in blood only. In our mathematical model, we thus assume homogeneous mixing of virus and cells throughout the body. It is important to note, however, that the characteristic decay profile in the study by Fischer et al. [12] could also be a result of differential trafficking of virus particles and HIV-1-infected subpopulations of cells between the blood and lymphoid tissue. Furthermore, it has also been suggested that the virion clearance rate from the blood corresponds to a virion efflux to other organs where the virus is ultimately cleared [32]. We assumed the death rates of cells during the intracellular eclipse phase (*I*_1_ to *I*_5_) to be the same as the death rate of CD4^+^ target cells. Some studies have suggested that infected cells in the eclipse phase could also be a target of cytotoxic T lymphocyte (CTL) killing and experience high death rates [24, 33–35]. The early steps of proviral transcription also remain elusive. It has been suggested that the decay of non-integrated viral DNA in infected cells could render them CD4^+^ target cells again [36–39]. The kinetics of HIV-1 DNA indeed show a small drop early after start of cART (Figure 2 and ref. [40]). However, we have excluded this effect for simplicity. Ultimately, the mechanisms of viral latency in HIV-1 remain a matter of debate [41]. In our model, we assumed that after proviral insertion some cells fail to increase viral RNA transcription and become latently infected cells. Latency could also result from infection of resting CD4^+^ T cells or de-activation of activated CD4^+^ T cells. We have not included the latter two mechanisms in our model as the data would not allow us to distinguish between them.

The complexity of the HIV-1 life cycle and its mathematical representation prevents the identification of a ‘true’ underlying model. We made several simplifying assumptions in our default model but we also studied a series of alternative models and found that some of those models also fit the data well. The limited number of patients prevents a more thorough analysis of the data. We also used the least-squares method to fit the model to the data and did not consider maximum likelihood approaches [42], values below the limit of detection or nonlinear mixed-effect models [43].

It remains to be determined how well the parameter estimates that were obtained during the chronic phase of infection represent the situation of acute HIV-1 infection. It is re-assuring that the simulated virus dynamics of acute infection show a peak around three weeks after infection which is in agreement with observations in patients [44,45]. Nevertheless, differences in immune activation during acute infection are likely to result in different proportions of cells becoming latent upon infection, and different activation rates of latently infected cells.

We found the HIV-1 burst size *in vivo* to be large, corroborating previous estimates from Chen et al. [46] who found the average burst size in SIV-infected rhesus macaques to be between 4.0 *×* 10^4^ and 5.5 *×* 10^4^. This is substantially higher than other estimates that were in the range of 10^3^ virions per cell [47, 48] and suggests that the number of virus-producing cells must be lower than previously anticipated. Measurements of extracellular virion-enclosed HIV-1 RNA (*Extra*) in the study by Fischer et al. [12] indeed suggest that the number of productively infected cells in peripheral blood is small which is also reflected in our model fits. In contrast to other studies that assume the viral production rate in longer lived persistently infected cells to be lower than in activated, virus-producing cells [49], we consider the viral production rates to be the same in both cell types. Instead, persistently infected cells can have occasional transcriptional bursts from *Mid* to *High* so they can release virus particles before reverting back to a lower transcriptional state or dying.

Our simulations of the development of different pools of HIV-1-infected cells are in good agreement with observations in patients. We find that the total number of HIV-1 DNA positive cells rapidly build up during the acute stage of infection. A very similar expansion was found in a recent study that measured the total number of HIV proviruses in PBMCs during the first weeks of HIV infection [50]. Also, our predicted ratio of the number of HIV-1 DNA positive cells during acute and chronic infection is in the same range as previously reported [31, 40]. The study by Murray et al. [40] further suggested that the level of HIV DNA continuously increases with duration of infection, reaching its 50% level at two years after infection. This contradicts earlier findings of stable levels of HIV-1 DNA positive PBMCs during the natural course of infection [51]. Our model predicts that the number of HIV-1 DNA positive PBMCs increases slowly during the first years of chronic infection and reaches its 50% level at 451 days after infection, corroborating the findings by Murray et al. [40].

An important question that remains is how many of HIV-1 DNA positive cells are latently or defectively infected. We found that the fraction of cells becoming defectively infected is surprisingly high. On the one hand, this could be a result of the assumption that HIV-1 DNA positive cells without viral RNA transcription remain silent. Some of these cells could actually be activated and start to produce UsRNA at low levels, i.e., become cells of the latent class *L*_1_. Eriksson et al. [52] measured a 300-fold difference between the number of latently infected cells as measured with a viral outgrowth assay and the total number of HIV-1 DNA positive resting CD4^+^ T cells. However, Ho et al. [53] showed a substantial fraction of noninduced proviruses in cells that have been stimulated in a viral outgrowth assay are replication-competent. They found that that the frequency of intact noninduced proviruses was at least 60-fold higher than the frequency of proviruses induced in a viral outgrowth assay. The median frequency of cells with intact noninduced proviruses per HIV-1 DNA positive resting CD4^+^ T cells was estimated at 3.7% [53]. In our simulation, the fraction of latently infected cells (*L*_1_ + *L*_2_) in all HIV-1 DNA positive cells (*DNA*^+^) is 1.8% (140.4/7608) during chronic infection. The striking correspondence of these numbers suggests that our mathematical model realistically describes the dynamics of the latent reservoir. Since the subpopulation of *L*_1_ is much larger than *L*_2_, the majority of latently infected cells consist of PBMCs that contain solely HIV-1 UsRNA (*Low)*, indicating that this transcriptional subclass is a good marker for viral latency.

This study provides an important step towards a more quantitative understanding of the dynamics of HIV-1 *in vivo*, in particular of the generation and maintenance of latently infected cells. A better understanding of the number of latently infected cells during acute infection is crucial for evaluating and predicting the outcome of early treatment and eradication strategies. Early cART treatment has been suggested to facilitate long-term control of HIV-1 [54] and studies have shown that it results in lower viral load levels during chronic infection [55]. Although the effects on viral load might only be transient [56], early treatment can prevent the expansion of viral cellular reservoirs in peripheral blood [31]. More recent strategies aim towards depletion of this reservoir [9], preferably during acute infection [57]. Predicting the chances of such eradication strategies critically depends on the ability to accurately quantify the pool of latently infected cells at various time points during HIV-1 infection. Our study shows that the latent reservoir becomes rapidly established during the first months of infection and represents a significant proportion (*>*1%) of all HIV-1 DNA positive PBMCs during chronic infection. In addition, our mathematical model realistically describes the dynamics of different HIV-1-infected subpopulations of cells which will be useful for projecting the effects of eradication strategies.

## Materials and Methods

### Patient data

We used previously published data from five chronically HIV-1-infected therapy naive patients that initiated cART using reverse transcriptase and protease inhibitors (patient numbers: 103, 104, 110, 111, 112) [12]. Plasma HIV-1 RNA (copies per ml) and CD4^+^ T cells (per µl) were measured at several time points during the first 48 weeks of cART. PBMCs were purified at weeks 0, 2, 4, 8, 12, 24 and 48 after the start of cART as described in Fischer et al. [58]. Serial dilution of PBMCs and patient matched PCR quantification of HIV-1 RNA species and DNA was performed as described elsewhere in detail [12, 13, 59, 60]. The freeze-thaw nuclease digestion method to differentiate between intracellular and virion encapsidated HIV-1 RNA has also been previously described in detail [4, 31]. HIV-1 RNA or DNA positive cell fractions measured as cells per 10^6^ PBMCs were converted to number of cells per ml of blood by multiplying with the number of PBMCs per ml. This ultimately lead to the stratification of cells to five (partially overlapping) subclasses [12]:

- *DNA*^+^: PBMCs containing HIV-1 DNA
- *Low*: PBMCs containing solely HIV-1 UsRNA
- *Mid*: PBMCs containing only HIV-1 MsRNA-tatrev or MsRNA-nef
- *High*: PBMCs containing elevated levels of both HIV-1 MsRNA-tatrev and MsRNA-nef
- *Extra*: PBMCs carrying virion-enclosed HIV-1 RNA

For the subclass *DNA*^+^, we make the assumption that there is only one proviral DNA copy per infected cell [61].

### Mathematical model

We devised a new virus dynamics model (Figure 3) which is adapted from previously published models [19, 24, 25, 30]. The various subpopulations of infected cells were stratified according to their HIV-1 DNA and RNA content. The model can be described by the following set of ordinary differential equations (ODEs):

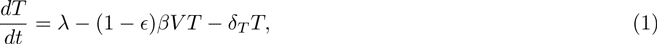

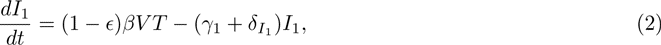

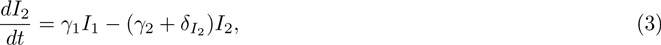

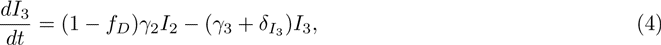

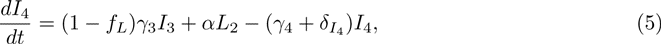

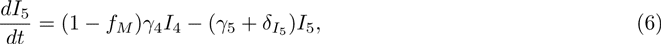

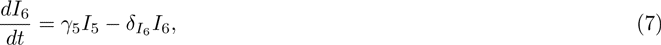

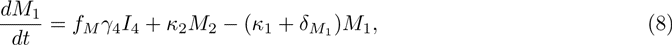

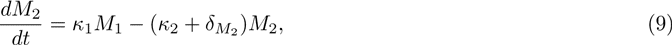

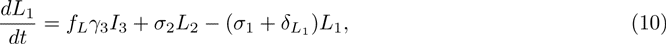

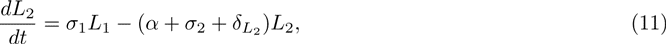

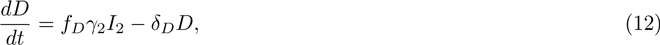

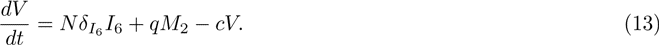

CD4^+^ target cells, *T*, are produced at rate *λ* and can become infected by virus particles, *V*, at rate *β*. *E* denotes treatment efficacy, where *E* = 0 before the start of antiretroviral therapy. Newly infected cells move through the intracellular eclipse phase, where *I*_1_ denotes the stage of reverse transcription, *I*_2_ the stage of proviral integration, and *I*_3_ to *I*_5_ subsequent stages with increasing transcriptional activity. After the intracellular eclipse phase, activated, virus-producing cells, *I*_6_, start to release free virus particles with a total viral burst size *N*. Some of the cells during the intracellular eclipse phase can become defectively infected cells, *D*, latently infected cells, *L*_1_, or persistently infected cells, *M*_1_. While we assume that defectively infected cells remain transcriptionally silent, both latently and persistently infected cells can exhibit transcriptional bursts that rise their transcriptional profile from *Low* to *Mid* and *Mid* to *High*, respectively. Latently infected cells in an elevated transcriptional state *L*_2_ can become activated at rate *α* or move back to the lower transcriptional state *L*_1_. Similarly, persistently infected cells that are highly transcriptionally active *M*_2_ can release free virus particles at rate *q* before they revert to a state of lower transcriptional activity or die. *δ_i_* and *c* describe cell death and viral clearance rates, respectively.

Due to the complexity of the full model, we make a number of simplifying assumptions. First, we assumed several of the cell death rates to be the same: the death rates of virus-producing cells (δ_*M*_2__ = δ*I*_6_), the death rates of defectively and latently infected cells (*δ*_*D* = *δ*_*L*_1__ = *δ*_*L*_2___) and the death rates of other infected cells and target cells (*δ*_*T*_ = *δ*_*I*_1__ = *δ*_*I*_2__ = *δ*_*I*_3__ = *δ*_*I*_4__ = *δ*_*I*_5__ = *δ*_*M*_1__). Second, the viral production rates in both virus-producing cells (*I*_6_ and *M*_2_) are kept the same, i.e., *Nδ*_*I*_6__ = *q*. Note, however, that persistently infected cells (*M*_2_) have a lower burst size than activated, virus-producing cells (*I*_6_) because they can revert to a non-productive state (*M*_1_). The default model described above is compared to a number of alternative models with different assumptions of the viral life cycle (*Text S1)*.

### Model fitting

The default model consists of 22 parameters of which 10 are fixed to previously used values from the literature (Table 1). The remaining 12 parameters were constrained and we used the geometric mean of the restricted range as starting values. The set of ODEs were solved numerically in the R software environment for statistical computing [62] using the function *ode* from the package *deSolve* [63]. We assumed that the chronic state of infection is reached after 1000 days (about three years), set *E* = 0.85 [22] and integrated the system during the time on cART (336 days).

The concentration of free virus *V* was measured directly but several of the infected cell populations contribute to the different subclasses of PBMCs (Figure 3): DNA^+^ = *I*_2_ + *I*_3_ + *I*_4_ + *I*_5_ + *I*_6_ + *M*_1_ + *M*_2_ + *L*_1_ + *L*_2_ + *D*, Low = *I*_1_ + *I*_3_ + *L*_1_, Mid = *I*_4_ + *L*_2_ + *M*_1_, High = *I*_5_ + *I*_6_ + *M*_2_ and Extra = *I*_6_ + *M*_2_. We further assume that target cells, *T*, correspond to a fraction, *s*, of all CD4^+^ T cells. Parameters were estimated by fitting the model to the data of each patient individually and minimizing the sum of squared residuals (SSR) between the prediction of the model and the data (taking the natural logarithm). All data points were weighted equally. However, the higher number of data points for free virus compared to cellular subclasses (e.g., *Extra*) forced the model to fit the virus concentration better than the other variables. We used the minimization algorithm by Nelder & Mead [64] that is implemented in the function *optim* and the *parallel* package for parallel computation. Parameter estimates are presented as geometric means including the ranges over all five patients. Code files can be obtained freely upon request from the corresponding author.

## Supporting Information

### Text S1

This file contains the calculation of the fate of cells after infection, the definition and results of the alternative models, the fits of the default model to the data from the four other patients, and the calculation of HIV-1 DNA positive cells from the study by Schmid et al. [31].

## Acknowledgments

CLA is funded by an Ambizione grant from the Swiss National Science Foundation (SNSF, grant 136737). Furthermore, the clinical and laboratory based work was supported by the Novartis Foundation (grant 02A03), Hartmann Müller Stiftung (grant 898), the Hermann Klaus Stiftung, an unrestricted educational grant by Abbott Inc. (grant SWIT-02-002), the Roche Research Foundation (grant no. 281-2005), the SNSF (grant 112670) and the University of Zurich’s Clinical Research Priority Program (CRPP) “Viral infectious diseases: Zurich Primary HIV Infection Study” (to HFG). ASP was supported by the National Institutes of Health (NIH, grant AI028433, OD011095, AI067854 (Center for HIV Vaccine Immunology) and AI100645 (Center for HIV Vaccine Immunology–Immunogen Discovery).

We would like to thank Roland Regoes for helpful discussions, the patients for participating in the study and Roland Hafner, Barbara Niederöst, Martina Ackerman, Viktor von Wyl, Philipp Kaiser and Rainer Weber for help in generating and analyzing the data. Part of this work was done during a research visit of CLA to the Los Alamos National Laboratory (LANL).

We want to remember our dear colleague and friend Marek Fischer, PhD who died in December 2010, without whom this work would not have been possible.

## Author Contributions

CLA designed the study together with HFG and BJ. CLA developed and performed the mathematical modeling, analyzed the results and wrote the manuscript. BJ and HFG provided data and analyzed the results. ASP contributed to the mathematical modeling and analyzed the results. All authors critically revised the manuscript and read and approved the final version.

